# Targeted sequencing of candidate genes of dyslipidemia in Punjabi Sikhs: Population-specific rare variants in *GCKR* promote ectopic fat deposition

**DOI:** 10.1101/526350

**Authors:** Dharambir K. Sanghera, Ruth Hopkins, Megan W. Malone-Perez, Cynthia Bejar, Chengcheng Tan, Huda Mussa, Paul Whitby, Chinthapally V. Rao, KarMing A. Fung, Stan Lightfoot, J Kimble Frazer

## Abstract

Dyslipidemia is a well-established risk factor for cardiovascular diseases. Although, advances in genome-wide technologies have enabled the discovery of hundreds of genes associated with blood lipid phenotypes, most of the heritability remains unexplained. Here we performed targeted resequencing of 13 bona fide candidate genes of dyslipidemia to identify the underlying biological functions. We sequenced 940 Sikh subjects with extreme serum levels of hypertriglyceridemia (HTG) and 2,355 subjects were used for replication studies; all 3,295 participants were part of the Asian Indians Diabetic Heart Study. Gene-centric analysis revealed a burden of variants for increasing HTG risk in *GCKR* (p=2.1×10^−5^), *LPL* (p=1.6×10^−3^) and *MLXIPL* (p=1.6×10^−2^) genes. Of these, three missense and damaging variants within *GCKR* were further examined for functional consequences *in vivo* using a transgenic zebrafish model. All three mutations were South Asian population-specific and were largely absent in other multiethnic populations of the Exome Aggregation Consortium. We built different transgenic models of human *GCKR* with and without mutations and analyzed the effects of dietary changes *in vivo*. Despite the short-term feeding, profound phenotypic changes were apparent in hepatocyte histology and fat deposition associated with increased expression of GCKR in response to a high fat diet (HFD). Liver histology of the *GCKR*^mut^ showed severe fatty metamorphosis which correlated with ~7 fold increase in the mRNA expression in the *GCKR*^mut^ fish even in the absence of a high fat diet. These findings suggest that functionally disruptive *GCKR* variants not only increase the risk of HTG but may enhance ectopic lipid/fat storage defects in the absence of obesity and HFD. To our knowledge, this is the first transgenic zebrafish model of a putative human disease gene built to accurately assess the influence of rare genetic changes and their phenotypic consequences *in vivo*.

## Introduction

Dyslipidemia is a well-established risk factor for cardiovascular disease and a principal cause of mortality in individuals with type 2 diabetes (T2D). Circulating blood lipid phenotypes are heritable risk factors for the development of atherosclerosis, and their measurements are used clinically to predict future coronary artery disease (CAD) risk and therapy for primary prevention^1, 2^. Epidemiological studies suggest that elevated serum triglyceride (TG) concentration is a strong independent risk factor for CAD^3,4,5^. There is an inverse correlation between serum TG and serum high-density cholesterol (HDL-C) that is associated with an increased risk of cardiovascular dysfunction, despite the level of low density cholesterol (LDL-C) being normal. This combination of lipid alterations is defined as atherogenic dyslipidemia, which is a significant risk factor for the development of CAD^6^. Lowering of LDL-C has been the major focus in CAD prevention following treatment with HMG-CoA reductase inhibitors (statins). However, the mortality rate of CAD remains elevated particularly in patients with T2D and insulin resistance, and reasons for their discordant effects in diabetics remain unknown^7^.

Family and twin studies have shown that TG and lipoprotein levels aggregate in families^8^. Relatives of individuals with hyperlipidemia/dyslipidemia will have a 2.5- to 7-fold increase in risk of death due to premature CAD compared to relatives of control individuals^9, 10^. The principal lipid alterations observed in these patients include high TG and low HDL-C. The Cincinnati Lipid Research Clinic Family Study showed that low HDL-C and high TG occur conjointly and are transmitted across generations as a “combined phenotype” or “conjoint trait”^11^. Genome-wide association studies (GWAS) and meta-analyses studies on multiethnic populations including Punjabi Sikhs have uncovered more than 200 genetic loci associated with circulating blood lipid phenotypes^12–15^. However, despite the high clinical heritability (50-80%) of many of the lipid traits^16^, these and several other studies have only explained up to 10% of heritability in these genes. To identify putative functional with larger effects, in this study, we have performed targeted sequencing of 13 bona fide candidate gene regions (~2.9 Mb) **(Table 1S/Figure 1S)** on 940 Sikh individuals [572 cases with high serum triglycerides (TG) (~ 95th percentile for their age and gender), and 368 controls with low TG (below the 20th percentile for their age and gender)], using subjects from the Asian Indians Diabetic Heart Study (AIDHS)/Sikh Diabetes Study (SDS)^17–19^.

## Materials and Methods

### Study Subjects of Discovery (sequencing) cohort

Genomic DNA samples of individuals including HTG cases (TG>150 mg/dl) and healthy controls with TG (<100 mg/dl) were sequenced with custom Nimblegen probes designed for target resequencing of 13 confirmed candidate genes for diabetic dyslipidemia in Sikhs. Diagnosis of T2D was confirmed by scrutinizing medical records for symptoms, use of medication, and measuring fasting glucose levels following the guidelines of the American Diabetes Association^20^. The diagnosis for normo-glycemic controls was based on a fasting glycemia <110 mg/dL or a 2-h glucose <140 mg/dL. CAD was assigned when there was a documented prior diagnosis of heart disease, electrocardiographic evidence of angina pain, coronary angiographic evidence of severe (>50%) stenosis, or echocardiographic evidence of myocardial infarction. HTG is broadly defined as fasting serum TG concentrations above the ninety-fifth percentile^21^ and was classified as mild HTG (150–399 mg/dL), high HTG (400–875 mg/dL), and severe HTG (>875 mg/dL).

The non-HTG control participants were recruited from the same Punjabi Sikh community and from the same geographic location as the HTG participants. They were selected on the basis of a fasting glycemia <100.8 mg/dL or a 2h glucose <141.0 mg/dL. BMI was calculated as weight (kg)/[height (m)^2^]. Education, socio-economic status, dietary and physical activity data were recorded. Smoking information was collected regarding past smoking, current smoking status, length of time, and number of cigarettes smoked /day. The vast majority of Sikhs were non-smokers, details are described elsewhere^19^. The individuals on lipid lowering medication are excluded from this cohort. All participants provided a written informed consent for investigations. The study was reviewed and approved by the University of Oklahoma Health Sciences Center’s Institutional Review Board, as well as the Human Subject Protection Committees at the participating hospitals and institutes in India. Metabolic estimations of fasting serum lipids [total cholesterol, LDL-C, HDL-C, and TG] were quantified by using standard enzymatic methods (Roche, Basel, Switzerland) as previously described^19^.

In this study, we only included individuals who self-reported having no South Indian admixture and exclusively belonged to the North Indian Punjabi Sikh community, and who reported that all four of their grandparents were of North Indian origin and spoke the Punjabi language. Excluded were individuals of South, East, or Central Indian origin; those of non-Sikh/non-Punjabi origin; those with rare forms of lipid disorders including very low serum TG (abetalipoprotenemia, homozygous hypobetalipoprotenemia, familial combined hypobetalipoproteinemia), or severe HTG (extremely high serum TG >1,000 mg/dL); those with familial chylomicronemia, hemochromatosis, or pancreatitis; those on lipid lowering medication; and those with excessive alcohol intake (>400 mL/day). About 50% of HTG patients had T2D and ~9% had CAD. Clinical characteristics of discovery-(resequencing) and replication cohort are summarized in **Table 1**.

**Table 1.** Phenotypic attributes of discovery (sequencing) and replication cohorts as part of Asian Indian Diabetic Heart Study (AIDHS)

| Traits | Discovery cohort<br>(n=820) |  | Replication cohort<br>(n=1,769) |  |
| --- | --- | --- | --- | --- |
|  | HTG cases | Controls | T2D cases | Controls |
| N | 572 | 248 | 1,074 | 695 |
| Females (%) | 39 | 49 | 44 | 46 |
| T2D (%) | 61 | 53 | N/A | N/A |
| Age (years) | 52.0 ± 12.4 | 55.0 ± 10.6 | 55.0 ± 11.7 | 45.7 ± 14.5 |
| BMI (kg/m<sup>2</sup>) | 27.6 ± 4.3 | 25.7 ± 4.4 | 27.1 ± 4.8 | 26.0 ± 4.6 |
| TG (mg/dl) | 314.9 ± 134.6 | 78.0 ± 16.9 | 158.1 ± 79.1 | 132.2 ± 67.6 |
| TC (mg/dl) | 203.3 ± 52.3 | 167.2 ± 42.7 | 178.3 ± 46.3 | 184.6 ± 90.2 |
| HDL-C (mg/dl) | 41.1 ± 17.5 | 41.9 ± 14.0 | 40.7 ± 14.1 | 42.6 ± 12.8 |
| LDL-C (mg/dl) | 110.6 ± 43.2 | 106.6 ± 36.2 | 107.9 ± 38.3 | 114.9 ± 35.7 |

HTG- hypertriglyceridemia, T2D-type 2 diabetes, BMI-body mass index, TG-triglyceride, TC-total cholesterol, HDL-C high density lipoprotein cholesterol, LDL-C low density lipoprotein cholesterol.

### Target Sequencing

Target sequencing was performed at the Northwest Genomics Center in the department of Genome Sciences at the University of Washington through the RS&G Service sponsored by the National Heart, Lung, and Blood Institute of the National Institutes of Health.

### Library Production, Targeted Capture, Sequencing

Library construction and custom capture were automated (Perkin-Elmer Janus II) in a 96-well plate format. 1 ug of genomic DNA was subjected to a series of shotgun library construction steps, including fragmentation through acoustic sonication (Covaris), end-polishing and A-tailing, ligation of sequencing adaptors, and PCR amplification with 8 bp barcodes for multiplexing. Libraries underwent capture using the Roche/Nimblegen SeqCap EZ custom designed probe. Prior to sequencing, the library concentration was determined by triplicate qPCR and molecular weight distributions verified on the Agilent Bioanalyzer (consistently 150 ± 15 bp). Barcoded libraries were pooled using liquid handling robotics prior to clustering (Illumina cBot) and loading. Massively parallel sequencing-by-synthesis with fluorescently labeled, reversibly terminating nucleotides was carried out on the HiSeq sequencer.

### Read Processing

Our sequencing pipeline is a combined suite of Illumina software and other “industry standard” software packages (i.e., Genome Analysis ToolKit [GATK], Picard, BWA, SAMTools, and in-house custom scripts), and consists of base calling, alignment, local realignment, duplicate removal, quality recalibration, data merging, variant detection, genotyping and annotation. The overall processing pipeline consists of the following elements: (1) base calls generated in real-time on the HiSeq2500 instrument (RTA 1.13.48.0) (2) demultiplexed, unaligned BAM files produced by Picard ExtractIlluminaBarcodes and IlluminaBasecallsToSam and (3) BAM files aligned to a human reference using BWA (Burrows-Wheeler Aligner; v0.6.2). Read data from a flow cell lane is treated independently for alignment and QC purposes in instances where the merging of data from multiple lanes is required (e.g., for sample multiplexing). Read-pairs not mapping within ± 2 standard deviations of the average library size (~150 ± 15 bp for exomes) were removed. All aligned read data are subject to the following steps: (1) “duplicate removal” was performed, (i.e., the removal of reads with duplicate start positions; Picard MarkDuplicates; v1.70) (2) indel realignment was performed (GATK IndelRealigner; v1.6-11-g3b2fab9) resulting in improved base placement and lower false variant calls and (3) base qualities were recalibrated (GATK TableRecalibration; v1.6-11-g3b2fab9).

### Sequence Data Analysis QC

All sequence data underwent a QC protocol before they were released to the annotation group for further processing. This included an assessment of: (1) total reads; (2) library complexity—the ratio of unique reads to total reads mapped to target (DNA libraries exhibiting low complexity are not cost-effective to finish); (3) capture efficiency—the ratio of reads mapped to human versus reads mapped to target; (4) coverage distribution—80% at 20X required for completion; (5) capture uniformity; (6) raw error rates; (7) Transition/Transversion ratio (Ti/Tv)—typically ~3 for known sites and ~2.5 for novel sites; (8) distribution of known and novel variants relative to dbSNP—typically < 7% using dbSNP build 129 in samples of European ancestry^22^; (9) fingerprint concordance > 99%; (10) sample homozygosity and heterozygosity and (11) sample contamination validation. All QC metrics for both single-lane and merged data were reviewed by a sequence data analyst to identify data deviations from known or historical norms. Lanes/samples that failed QC were flagged in the system and could be re-queued for library prep (< 5% failure) or further sequencing (< 2% failure), depending upon the QC issue. Completion was defined as having > 80% of the target at >20X coverage.

### Variant Detection

Variant detection and genotyping were performed using the UnifiedGenotyper (UG) tool from GATK (v1.6-11-g3b2fab9). Variant data for each sample were formatted (variant call format [VCF]) as “raw” calls that contain individual genotype data for one or multiple samples, and flagged using the filtration walker (GATK) to mark sites that were of lower quality/false positives [e.g., low quality scores (Q50), allelic imbalance (ABHet 0.75), long homopolymer runs (HRun >3) and/or low quality by depth (QD <5)].

### Variant Annotation

We used an automated pipeline for annotation of variants derived from exome data, the SeattleSeq Annotation Server (http://gvs.gs.washington.edu/SeattleSeqAnnotation/). This publically accessible server returns annotations including dbSNP rsID (or whether the coding variant is novel), gene names and accession numbers, predicted functional effect (e.g., splice-site, nonsynonymous, missense, etc.), protein positions and amino-acid changes, PolyPhen predictions, conservation scores (e.g., PhastCons, GERP), ancestral allele, dbSNP allele frequencies, and known clinical associations. The annotation process has also been automated into our analysis pipeline to produce a standardized, formatted output (VCF-variant call format, described above).

### Replication studies, Population Characteristics, and SNP Genotyping

We replicated the association of three functional variants in an additional 2,355 individuals of Punjabi Sikh ancestry. These included 1,000 individuals from Sikh families and the remaining 1,355 were unrelated; all were part of the AIDHS/SDS as described previously^19, 23–25^. Recruitment and diagnostic details of the Sikh replication cohort are similar as described above for the discovery cohort. Clinical and demographical details of these cohorts are provided in **Table 1.** Genotyping for selected SNPs was performed using TaqMan pre-designed or TaqMan made-to-order SNP genotyping assays from Applied Biosystems Inc. (ABI, Foster City, USA) as described previously^26^. Genotyping reactions were performed on the QuantStudio 6 genetic analyzer using 2 uL of genomic DNA (10 ng/uL), following manufacturers’ instructions. For quality control, 8-10% replicative controls and negative controls were used in each 384 well plate to match the concordance. Genotyping call rate was 96% or more in all the SNPs studied.

### Functional Studies using Zebrafish (ZF) Model

To test the phenotypic effects of this and other novel variants *in vivo,* we created transgenic ZF (*Danio rerio*) models of the glucokinase regulatory protein (*GCKR)*^*mut*^, and *GCKR*^*wt*^ using the TAB-5 strain, a commonly used strain derived from two commonly used fish lines (Tubingen and AB). Heterozygous human carriers of this mutation exhibit HTG and high rates of T2D, thus we examined whether *GCKR*^*mut*^ induces features of this phenotype in ZF. To build our transgenic models, we employed the *Tol2* system, which mediates highly-efficient transgenesis^27^. We used a promoter construct that drives human *GCKR*^*mut*^ expression only in hepatocytes, while simultaneously labeling those cells fluorescently. To achieve hepatocyte-specific expression in *D. rerio*, we used the *D. rerio* liver fatty acid binding protein (L-FABP) promoter (courteously provided by Dr. Schlegel, University of Utah)^28^. To label *GCKR*^*mut*^ expressing *D. rerio* hepatocytes, we joined the cDNA for human *GCKR*^*mut*^ to the enhanced red fluorescent protein (mCherryFP), separated by a 2A peptide linker^27, 29^. The 2A linker is an auto-cleaving peptide, resulting in the *GCKR*^*mut*^ and mCherryFP proteins being expressed in a 1:1 stoichiometric ratio. Expression of mCherryFP by the ZF liver confirms expression of *GCKR*^*mut*^ hepatocytes **(Figure 1A-C)**, and also facilitates purification of *GCKR*^*mut*^-expressing cells by Fluorescence Activated Cell Sorting for downstream analyses. After building *GCKR*^*mut*^ transgenic ZF, we evaluated the *in vivo* metabolic consequences of these human *GCKR* mutations by feeding high fat diet to 5 day old larvae of wildtype (WT) and transgenic ZF with and without GCKR mutations.

**Figure 1 A-C:**
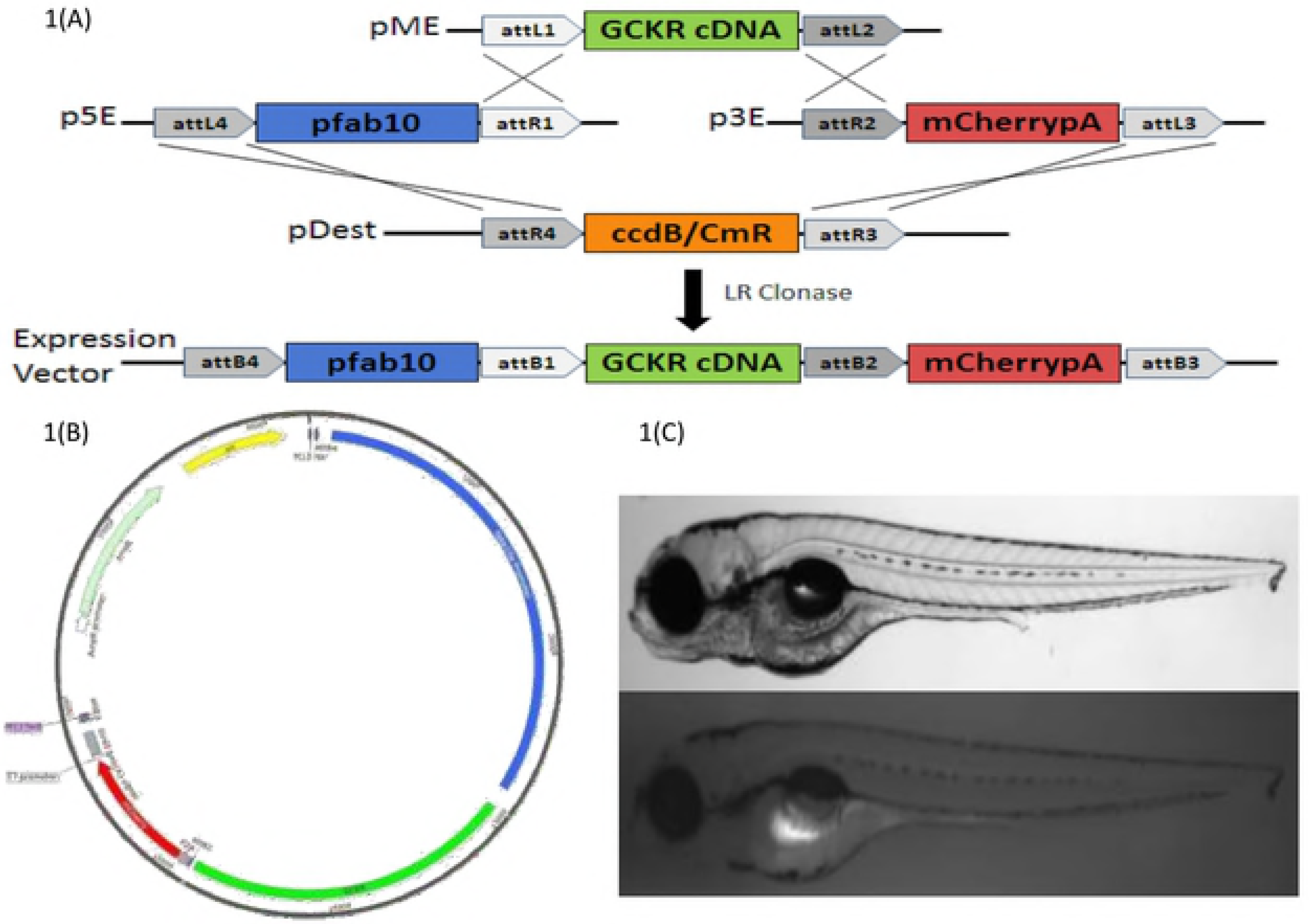
Transgenic ZF Model Construction. (A) Multisite Gateway-based plasmid construction using *Tol2* system; (B) Bacterial plasmid construct (p3E-2A-mCherrypA); (C) larvae (4 day post fertilization) of wild type and transgenic fish showing hepatic expression (fluorescence) after successful transgenesis of human *GCKR* with three disruptive mutations.

### Diet Experiments

For all feeding studies, 5-day post-fertilization (dpf) homozygous humanized *GCKR*-mutant or–WT transgenic zebrafish larvae were studied. For comparison to WT, we also studied 5 dpf larvae of Tab-5 fish (the parental line used to construct transgenic constructs). All WT and mutant fish were distributed in 3 liter tanks (20 fish per tank) and fed defined diets for 14 days. Animals were housed in the main aquarium of the ZF Animal Resource core facility at the University of Oklahoma and maintained on a 14-hour light, 10-hour dark cycle. Animals were anesthetized and killed by immersion in ice water^30^. For control studies, 5 dpf homozygous *GCKR* mutant (*GCKR*^mut^) or homozygous WT (*GCKR*^wt^) fish were reared on a conventional diet (commercial powder and newly-hatched *Artemia salina nauplii*), twice-daily for 14 days. The high fat diet (HFD) groups were fed with a special fish diet (with 24% fat, 43% protein, and 4% fiber from Purina Aqua Mix) thrice-daily for 14 days. Three larvae from each feeding group were euthanized and their livers were dissected using a dissection microscope and sent for transmission electronic microscopy at the Oklahoma Medical Research Foundation’s imaging core facility.

### Larvae tissue embedding and Hematoxylin and Eosin (H&E) staining

A Leica TP1020 tissue processor was used to process the tissue, following the manufacturer protocol. Briefly, tissue in 10% neutral formalin buffer (NBF) are moved into labelled tissue blocks. The tissue in the blocks are progressively dehydrated with increasing concentrations of ethanol, then in xylene and imbibed with paraffin liquid. Due to fragility of Zebrafish larvae, they were placed in biospecimen bags and 5 minutes in each step of processing was adapted. The paraffin imbibed tissue is taken out and embedded according to orientation as needed using a 10X dissection microscope. The formalin-fixed paraffin-embedded tissues were sectioned at desired thickness (4 μm) and mounted on positively charged slides. The slides were dried overnight at room temperature and incubated at 60°C for 45 minutes. The Hematoxylin and Eosin were purchased from Leica Biosystems, and staining was performed utilizing Leica ST5020 Automated Multistainer following the Hematoxylin-Eosin (HE) staining protocol at the SCC Tissue Pathology Shared Resource.

### Quantitative gene expression studies

Gene expression studies for quantifying GCKR mRNA was performed on ZF larvae fed with normal and HFD. Total RNA was isolated using the Absolutely RNA Mini Prep Kit (Agilent Technologies Inc., Santa Clara, CA), and was reverse transcribed using the iScript cDNA Synthesis Kit (Bio-Rad Laboratories), according to the manufacturers’ protocols. For the quantification of GCKR mRNA, quantitative PCR (qPCR) was performed using SsoAdvanced SYBR Green Supermix (Bio-Rad Laboratories, Hercules, CA). Real Time qPCR was then performed using QuantStudio 6 in conjunction with *GCKR* forward and reverse primers (Integrated DNA Technologies, Skokie, Illinois, USA) and Bio-Rad’s SYBR Green Supermix with ROX) (**Supplementary Table 4S**). Beta-actin was used as a normalizing control. Results were analyzed using ABI’s QuantStudio Real-Time PCR software (v.1.3).

### Bioinformatics and Statistical analysis

Missense variants were designated as damaging using the *in-silico* predictions generated by tools like PolyPhen^31^, SIFT^32^, BONGO^33^, LRT^34^, Mutation Taster^35^, and PolyPhen-2^36^. The variants with a score of four out of six defined by these algorithms were considered potentially damaging. Data quality for SNP genotyping was checked by establishing reproducibility of control DNA samples. Departure from HWE of common variants in controls was tested using the Pearson chi-square test.

### Gene-centric Association Analysis

For gene-centric analysis, we performed gene burden tests to jointly analyze multiple non-synonymous or other likely functional variants, including singleton variants by Combined Multivariate and Collapsing (CMC) method^37^, implemented in SVS v.2.0 (Golden Helix, Bozeman, MT, USA). This method applies collapsed rare variants in different MAF categories to evaluate the joint effect of common and rare variants. We also used the variance-component test within a random-effects model including the sequence kernel association test (SKAT)^38^, which tests for association by evaluating the distribution of genetic effects for a group of variants instead of aggregating variants.

### Single SNP Association Analysis

The genotype and allele frequencies in T2D cases were compared to those in control subjects using the chi-square test. Statistical evaluation of genetic effects on T2D risk used multivariate logistic regression analysis with adjustments for age, gender, and other covariates. Continuous traits with skewed sampling distributions (e.g., triglycerides or fasting glucose) were log-transformed before statistical analysis. However, for illustrative purposes, values were re-transformed into the original measurement scale. General mixed linear models were used to test the impact of genetic variants on transformed continuous traits using the variance-component test, adjusted for the random-effects of relatedness and fixed effects of age, gender, BMI and disease implemented in SVS v.2.0 (Golden Helix, Bozeman, MT, USA). Other significant covariates for each dependent trait were identified by Spearman’s correlation and step-wise multiple linear regression with an overall 5% level of significance using SPSS for Windows statistical package (version 18.0) (SPSS Inc., Chicago, USA). Mean values between cases and controls were compared by using an unpaired t test. To adjust for multiple testing, we used Bonferroni’s correction (0.05/number of tests performed).

## Results

Of a total of 2,709 individuals studied, target sequencing was performed on 940 subjects and 1,769 subjects were used for the replication studies. All these participants were part of the AIDHS/SDS^17–19^. Of the 940 sequenced samples, 820 passed the stringent QC based on multiple parameters and were used for further analysis. **Table 1S** describes details of the lipid candidate gene regions selected for targeted resequencing. A total of 2,361 high-quality variants were analyzed for their distribution and association with lipid-related traits, diabetes and other cardiometabolic risk factors **(Table 2S)**.

Our results revealed accumulation of several unknown rare (<1%) and less common variants (<10%) that were not found in any of the existing variant databases. For instance, our results of *GCKR* sequencing in Sikhs revealed clustering of 13 rare mutations and many of these were predicted to be damaging/ deleterious based on the *in-silico* prediction methods **(Figure 2 A)**. Gene-centric analysis for studying the aggregate effects of clustered variants within each gene, revealed a significant burden in *GCKR* (p=2.1×10^−5^) along with *LPL* (p=1.6×10^−3^) and *MLXIPL* (p=1.6×10^−2^), for increasing the risk for HTG **(Table 1s).**

**Figure 2A-E:**
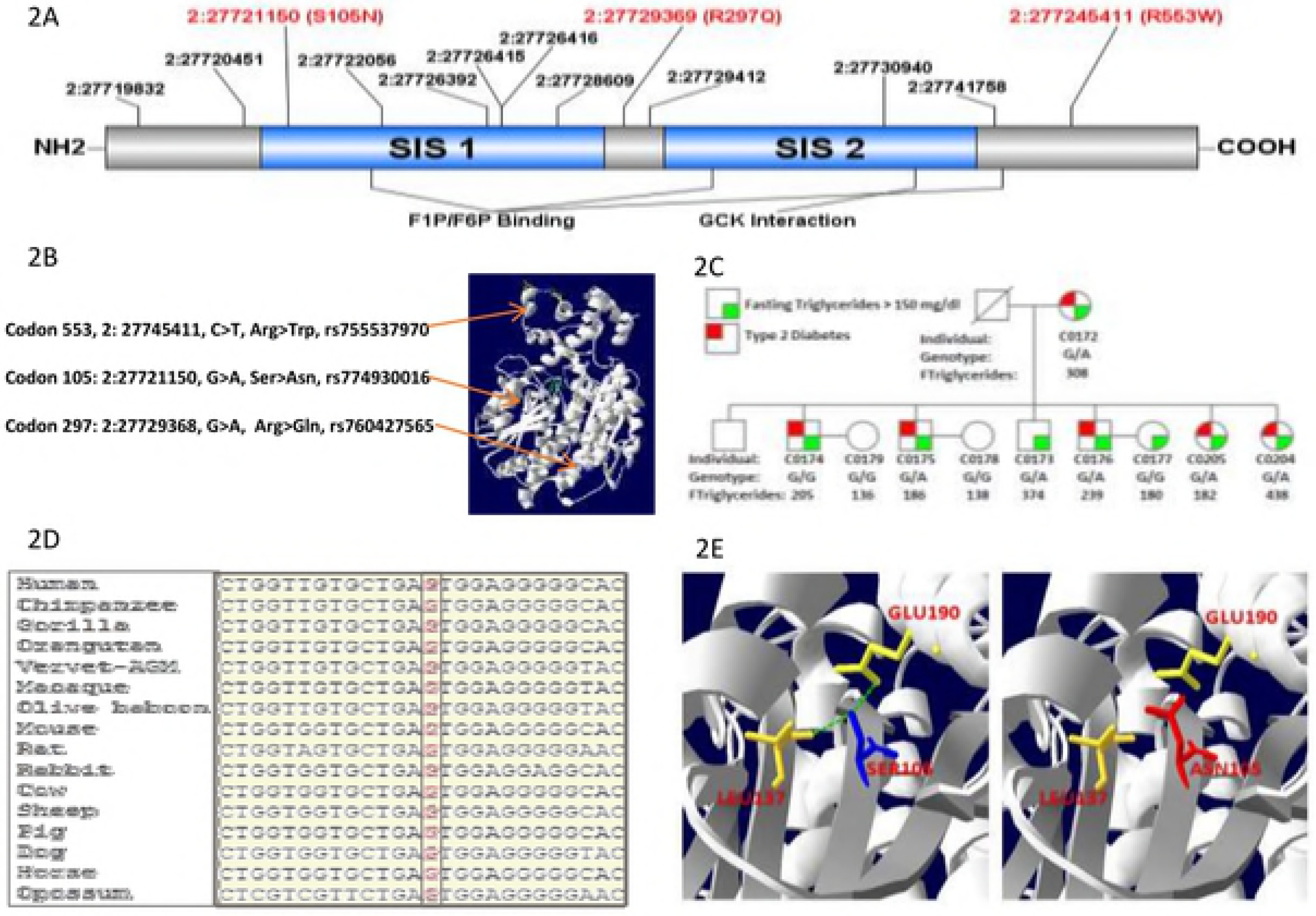
Population-specific rare functional variants. 2A. Target sequencing in the *GCKR* gene reveals three novel damaging mutations in SIS domains in Sikhs; 2B. Crystal structure of human GCKR showing mutant residues (S105N) mapping to SIS-1 FIP/F6P binding domain while R297Q is located between SIS-1 and SIS-2 domains, while R553W maps near the GCK interaction domain; 2C. Pedigree figure of a Sikh family showing overrepresentation of a rare damaging variant rs774930016 (S105N) segregating with T2D and hypertriglyceridemia; 2D. Sequence alignment of SIS-1 domain reveals absolute conservation of rs774930016 at position S105N of the GCKR gene across species; 2E. The wild-type residue (blue) forms hydrogen bonds with Glutamic Acid at position 190 and Leucine at position 137. However, the mutant residue (red) destabilize the folding of Fructose Binding domain by the loss of a hydrogen bond with Glutamic Acid 190 and Leucine at position 137.

The present study is further mainly focused on 3 population-specific rare variants identified in the *GCKR* gene **(Figure 2 B).**The first functional variant (S105N), located on the Sugar Isomerase domain −1 (SIS-1), was functionally disruptive and absent in Caucasians (n=33,356), Africans (n=5,197), Hispanic/ Latinos (5,788), and East Asians (n=4,324) in a large Exome Aggregation Consortium (ExAC) of multiethnic populations **(Table 2)**. Two additional rare functional variants (R297Q and R553W) were also confined to this Sikh population only, and most carriers had HTG **(Figure 2A-B, Table 2).** Two of these three disruptive missense variants (S105N near the fructose binding site and R553W near the GCK interaction domain) were highly conserved across species **(Figure 2D, Figure 3B1)**, while the mutant allele of R297Q variant was also found in cow and sheep in addition to its predominant presence in South Asians **(Figure 3A1)**.

**Table 2.** Carrier counts for three population-specific variants in the *GCKR* gene in AIDHS and multiethnic populations.

| Population | rs774930016 (S105N) | rs760427565 (R297Q) | rs755537970 (R553W) |
| --- | --- | --- | --- |
| European | 0/33356 | 1/33359 | 1/33358 |
| Latino | 0/5788 | 1/5787 | 0/5788 |
| African | 0/5197 | 0/5200 | 0/5197 |
| East Asian | 0/4324 | 0/4326 | 0/4327 |
| South Asian | 3/8250 | 11/8255 | 1/8255 |
| AIDHS (Sikhs) | 9/3132 | 18/3016 | 8/2950 |

Data of all non-Sikh populations are from Exome Aggregation Consortium (ExAC).

**Figure 3 A-B:**
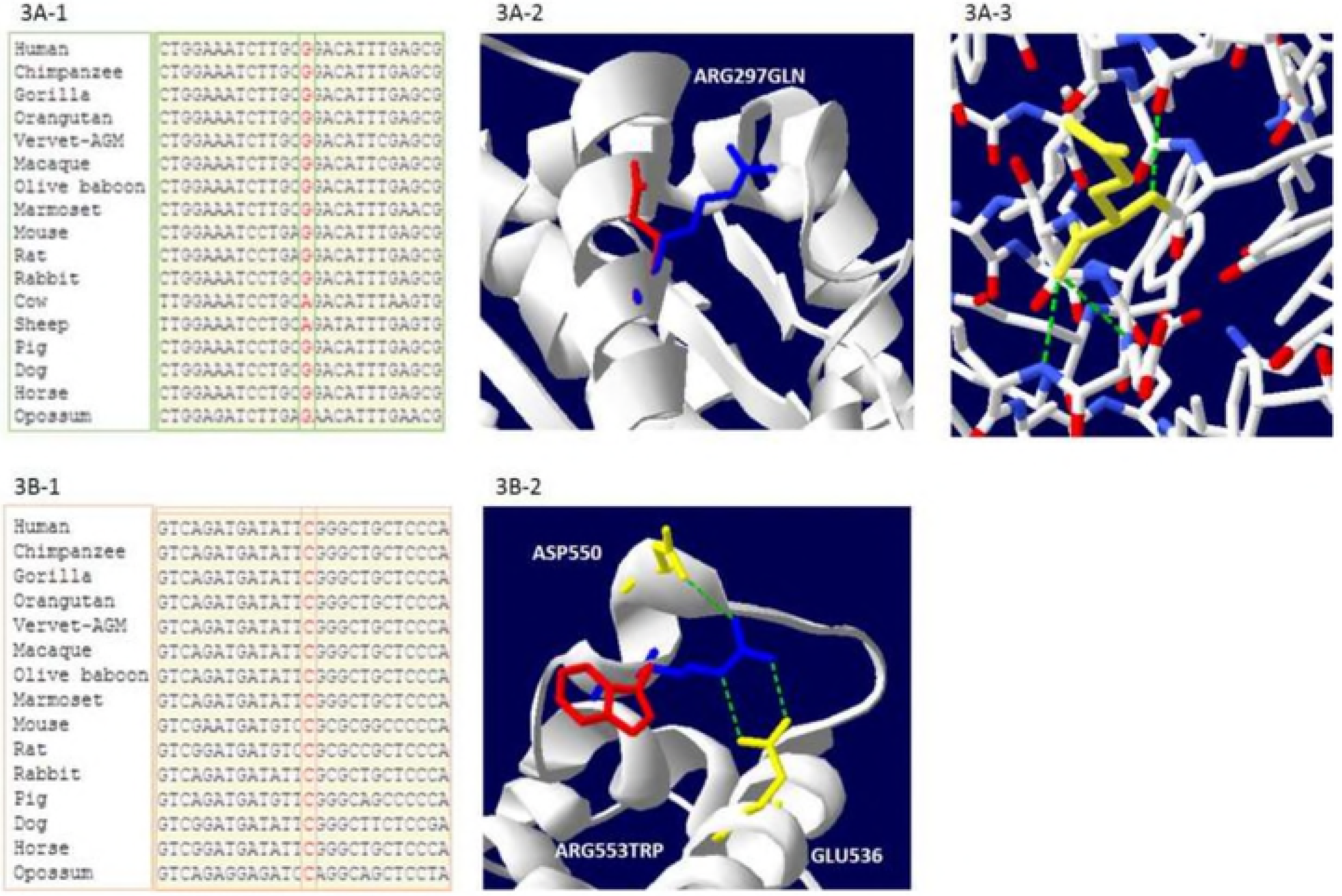
Population-specific rare functional variants of interest. 3A-1. Sequence alignment of 297Gln reveals partial conservation of ancestral ‘G’ allele at position Arg297 to Gln297 of the GCKR gene across species. 3A-2. Protein modeling of Arg297Gln showing the wild-type blue (Arg297) and mutant red (Gln297) residues. 3A-3. The wild-type Arg297 forms a salt bridge with glutamic acid at position 63 and glutamic acid at position 300, however, the new mutant residue is not in the correct position to make the same hydrogen bond as the original wild-type residue did. 3B. 3B-1. Sequence alignment reveals a complete conservation of wild type ‘C’ allele at position Arg553Trp of the GCKR gene across species. 3B-2. Protein modeling of Arg553Trp mutations in the GCKR gene. The wild-type residue (blue) forms a hydrogen bond with glutamic acid at position 536, and aspartic acid at position 550. The size difference between the wild-type and mutant residue makes it to where the new residue is not in the correct position to make the same hydrogen bond as the original wild-type residue. The wild-type residue was positively charged, the mutant residue is neutral. The mutant residue is more hydrophobic than the wild-type residue.

The three functionally tested damaging rare mutations in *GCKR* were at the fructose binding site and GCK binding site at or near the sugar isomerase (SIS-1-2) domains **(Figure 2A).** The disruptive allele at codon 105 is predicted to destabilize the folding of the fructose binding domain that results in the loss of a hydrogen bond between Serine (105) and Glutamine (190) **(Figure 2E)**. Interestingly, this variant was monomorphic in Europeans, East Asians, Africans and Latinos of the ExAC consortium, and only 3 of 8250 South Asians from Pakistan were carriers (genotype frequency 0.00036), whereas 9 out of 3132 Sikhs were carriers of this variant (0.0029). This variant co-segregated between heterozygous carriers, HTG- and T2D phenotypes in one Sikh family. Of these over 83% of carriers in this family had HTG (ranging from 148mg/dl to 530 mg/dl) and 75% of carriers were diabetic. Similarly, two more rare functional variants (R297Q and R553W) were confined to this population only and were with high TG in most individuals **(Supplementary Tables 3SB-C).**

We investigated single variant association of each rare variant with diabetes and quantitative risk phenotypes (e.g. fasting glucose, body mass index (BMI), total cholesterol, LDL-C, HDL-C and TG) in the discovery and replication cohorts. None of these variants showed any significant association with diabetes, fasting glucose or lipid traits (data not shown) except TG. As shown in **Table 3,** carriers of the S105N (rs774930016) variant had significantly increased serum TG levels (β 0.59 ± 0.17; p=7 × 10^−4^) after adjusting for age, gender and BMI. This association remained significant even after including T2D and family relatedness in the model (β 0.60 ± 0.17; p=4.32 × 10^−4^) in the replication cohort and in combined (discovery and replication) samples (β 0.55 ± 0.19; p=0.004). A similar but marginally significant association of R553W (rs755537970) variant, was observed in combined samples with triglycerides (β 0.51± 0.23; p=0.028). However, no significant association was observed in R297Q (rs760427565) with TG **(Table 3, Figure 6)**.

**Table 3.**
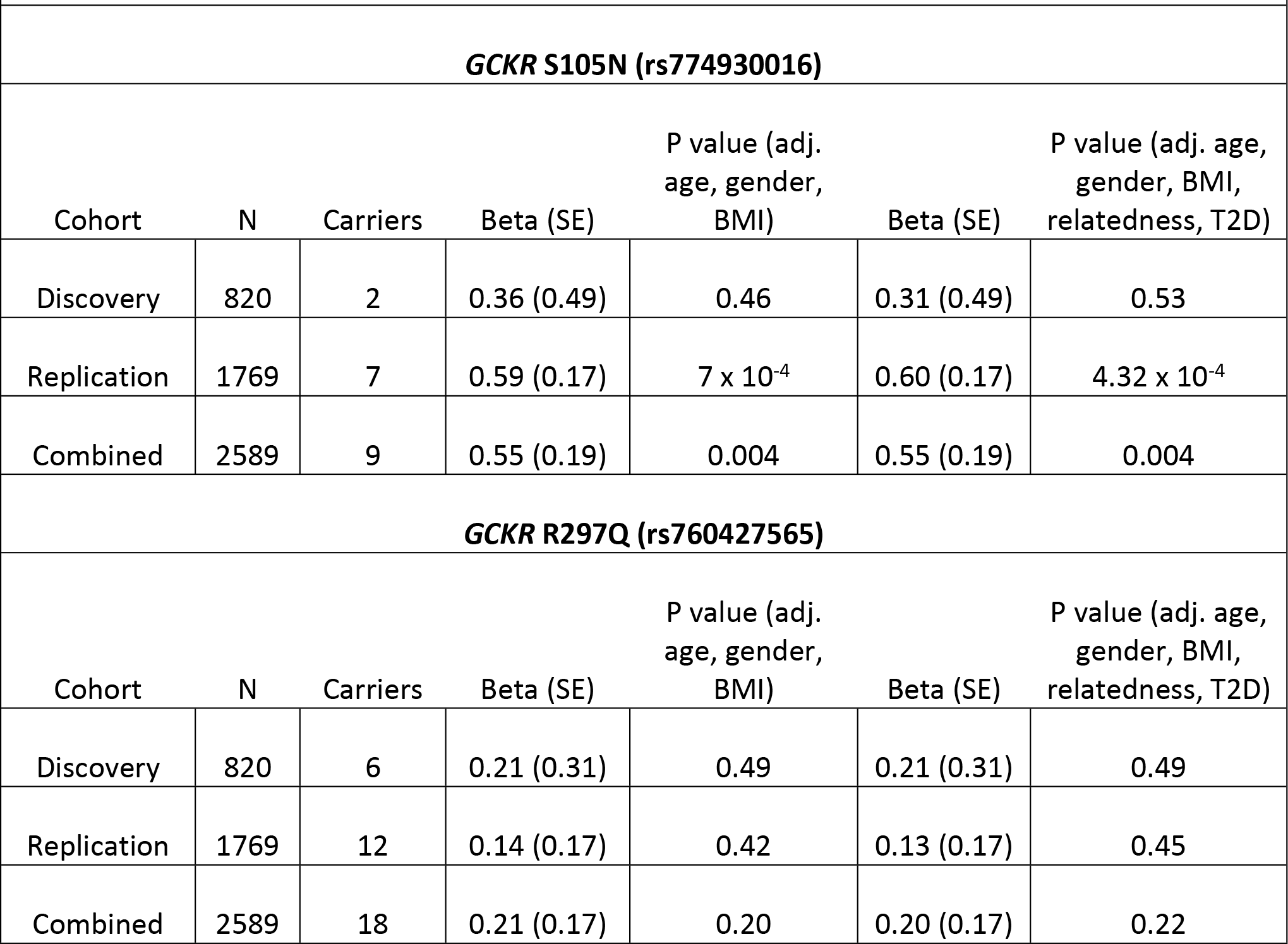
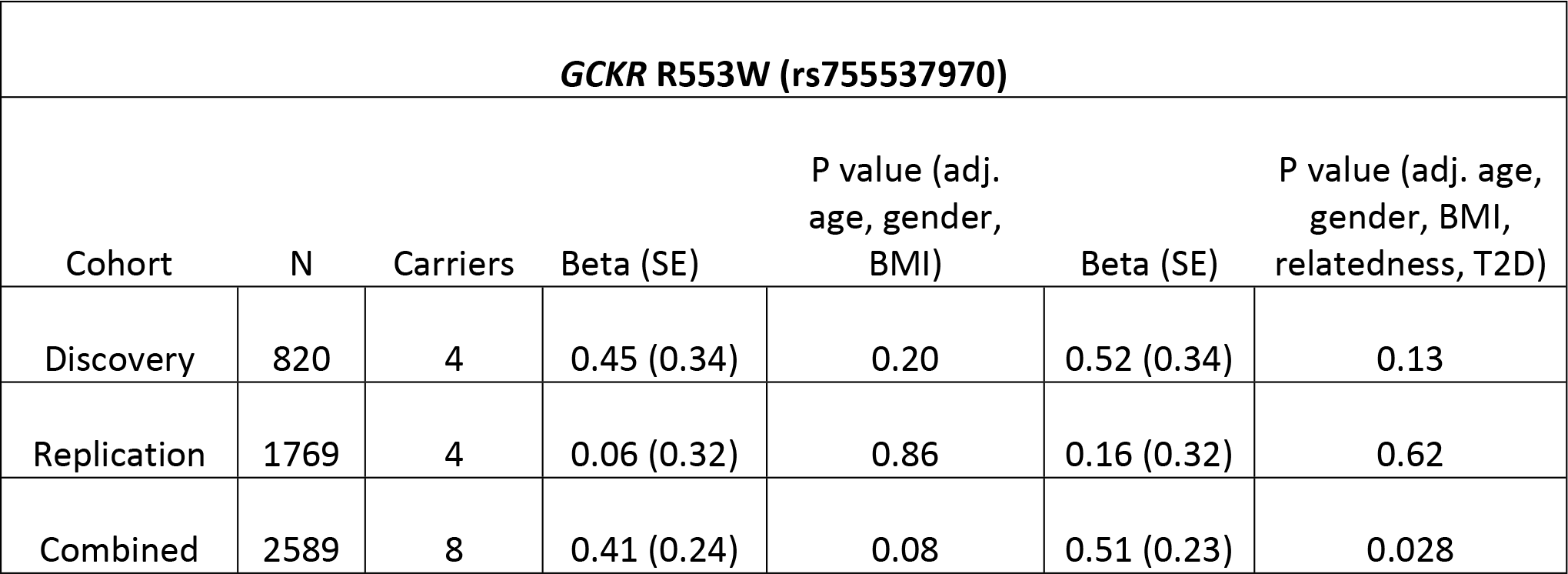
Multivariate linear regression analysis showing association of three population-specific rare *GCKR* variants with serum triglycerides in discovery and replication cohorts.

Based on the significant association of these variants with HTG, we next evaluated the functional consequences of three South Asian population-specific variants by designing a humanized GCKR ZF model. Expression of mCherryFP by the ZF liver confirms expression of *GCKR*^*mut*^ hepatocytes **(Fig 1A-C)**. The H&E liver images of TAB-5, transgenic *GCKR* ^wt^, and *GCKR* ^mut^ groups with normal diet and HFD are shown in **Figure 5A-C**. The fat deposition in liver hepatocytes of TAB-5 larvae was increased 3-4-fold in response to HFD. A similar increase in response to HFD was noticed in transgenic fish with wild type GCKR. However, the mutant transgenic fish exhibited a 3-fold increase in ectopic fat in hepatocytes with a normal diet, with 80% hepatocytes having fat deposition, while transgenic mutants on HFD had hepatocytes loaded with fat showing a marked degeneration of hepatocyte nuclei with possible steatosis **(Figure 5C).** Additional supplemental figures of WT and transgenic Zebrafish livers at 40X and 4x magnification are shown in **Figure 3S** and **Figure 4S**, respectively. In response to a HFD, mRNA expression of GCKR increased about two folds in normal TAB-5 compared to a normal diet. On the other hand, there was 7-fold increase in *GCKR* ^*mut*^ larvae even in the absence of HFD; whereas, the *GCKR* ^*mut*^ mRNA levels were restored to normal when fed on HFD **(Figure 4)**.

**Figure 4.**
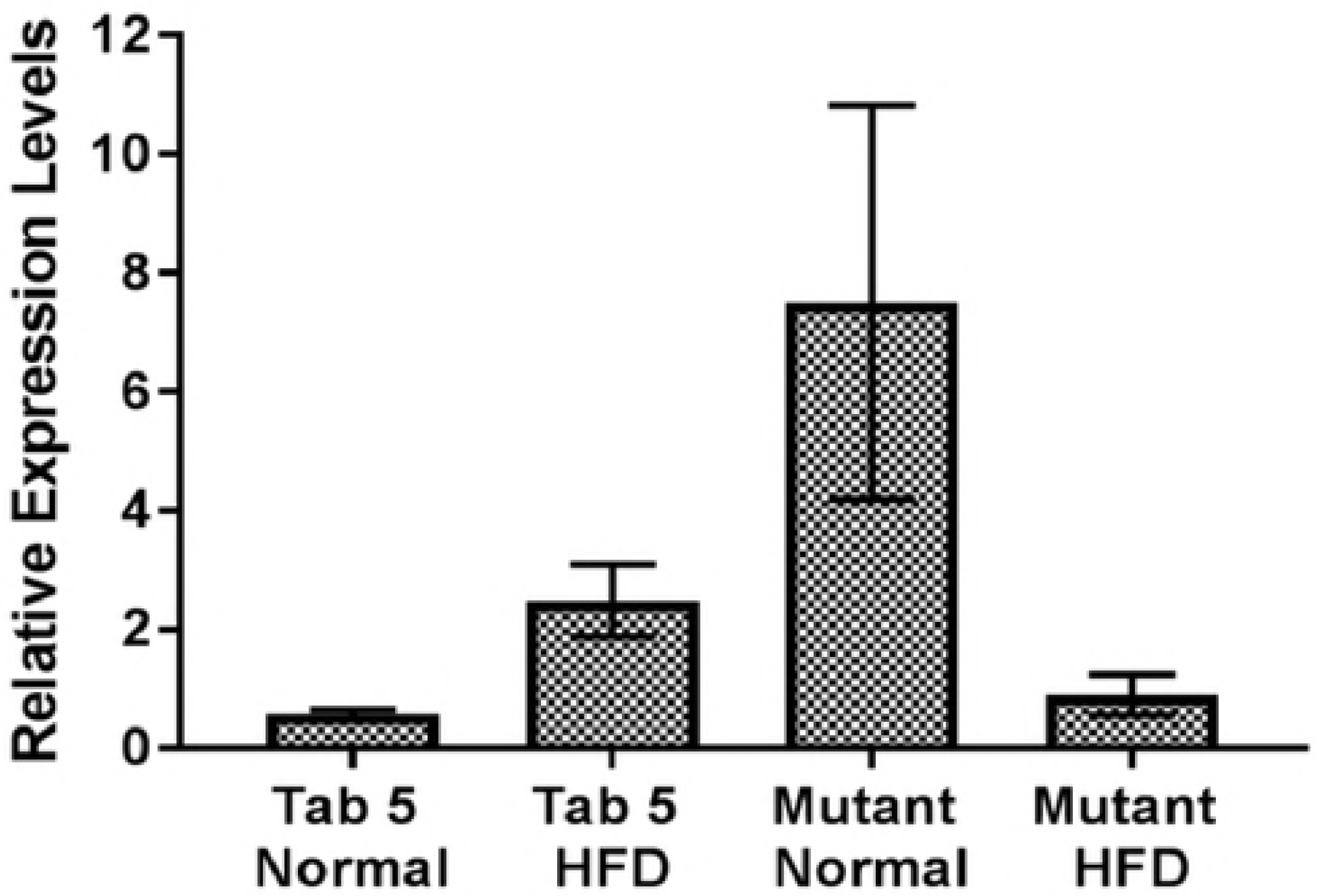
GCKR Relative Expression Levels. Graph summarizes the relative mRNA expression levels of GCKR normalized by beta actin in zebrafish larvae of wildtype (WT) Tab-5 and transgenic mutant fish fed on a normal and high fat diet (HFD). Data are presented as mean and standard error of mean using Tab-5 WT as reference. Significant differences in the relative expression of GCKR mRNA (denoted by an asterisk) were detected between Tab-5 (WT) and the transgenic mutant on a normal diet; and between transgenic mutants on a normal diet vs. HFD.

**Figure 5.**
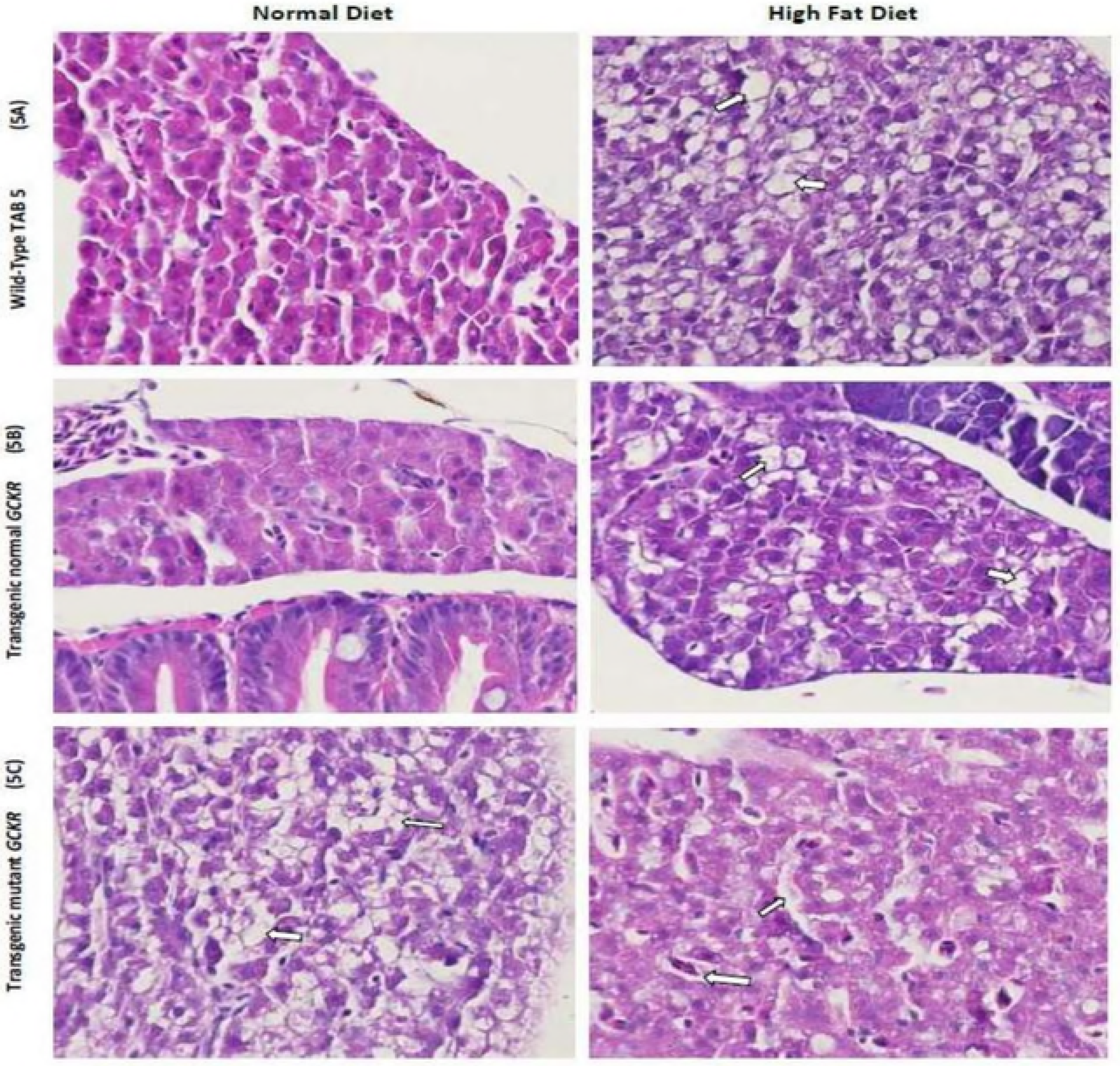
Zebrafish liver histology of Hematoxylin and Eosin stained sections showing cellular differences in normal and high fat feeding for 2 weeks. (5A). A wild-type Tab-5 liver exhibited normal hepatic cells, while the HFD liver was normal with scattered fat cells (white arrow) indicating 3+ fat without apoptosis. (5B) Transgenic fish with wild type GCKR showed normal hepatocytes in larvae fed a normal diet, and a normal liver with scattered fat cells (white arrow) in transgenic larvae fed a HFD with no apoptosis. (5C) Transgenic mutant GCKR fed with a normal diet showed a liver with fatty metamorphosis and scattered hepatic cell apoptosis (white arrow) with 4+ fat, and with approximately one apoptotic cell every hundred hepatic cells. Transgenic mutant GCKR fed with a HFD shows 4+ fat with severe cellular damage and higher fatty infiltration than the normal and transgenic normal fish. Most hepatic cells in mutant transgenic fish with and without a HFD contain vacuoles of fat and show severe disorganization of the hepatic structure and hepatocyte nuclei with possible steatosis.

**Figure 6.**
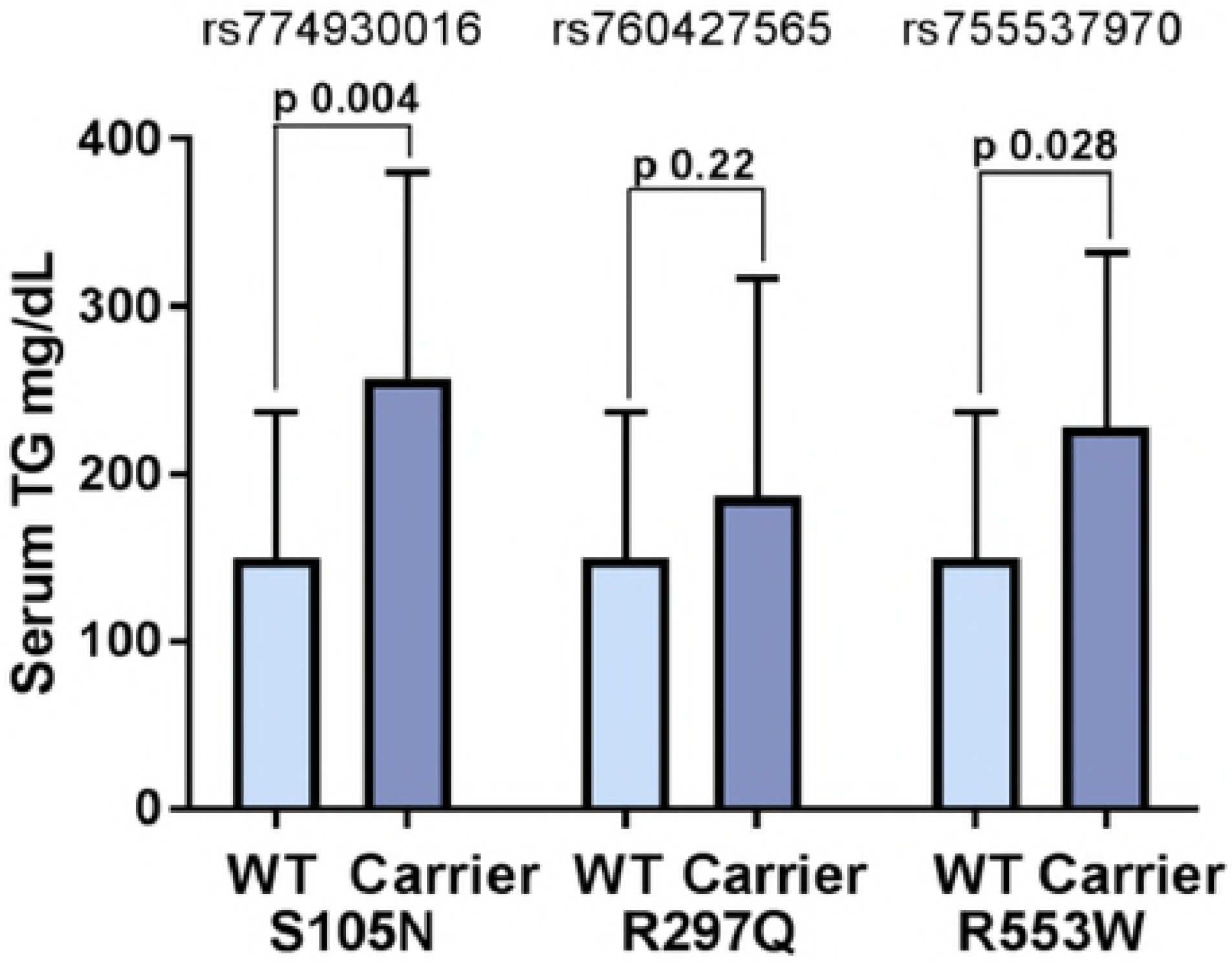
Serum Triglyceride mean distribution among carriers and non-carriers of missense variants. Bar graph shows the differences in the distribution of mean serum triglycerides among carriers and non-carriers of missense mutations (S105N, R297Q, and R553W) represented by rs774930016, rs760427565, rs755537970 variants, respectively. Data are shown in mean and standard deviation of means. *P* values are derived from the general mixed linear models used to test the impact of genetic variants on transformed continuous trait (TG) using the variance-component test adjusted for the random-effects of relatedness and fixed effects of age, gender, BMI and type 2 diabetes. Only association of S105N would remain significant after applying Bonferroni correction.

## Discussion

In this investigation, we have attempted to identify functional variants by resequencing 13 known candidate genes of dyslipidemia using an endogamous population of Punjabi Sikhs known to have a high risk for cardiovascular diseases^18, 19, 25, 39^. Despite considerable success of GWAS, whole-genome and exome sequencing, including studies from our group^17, 40–42^, the genetic mechanisms that predispose people to metabolic and cardiovascular disease risk factors remain poorly understood. The degree of clinical heterogeneity existing in the CAD or cardiometabolic phenotypes imposes serious limitations in our ability to effectively measure genetic risk, environmental exposure, and their interactions. Additionally, most post-GWA studies on candidate gene sequencing have predominantly been focused on European populations which provide limited information on the usefulness of variants in populations of non-European ancestry. Moreover, the post-GWAS exome arrays could capture the majority of low-frequency variants in European populations only when the sample size exceeds >300,000^43^. However, such studies in other disparate populations are sparse. The current investigation in family and population based samples from the AIDHS/SDS is an effort to identify missing heritability associated with GWAS-driven loci of dyslipidemia, specifically the HTG by using candidate gene resequencing.

As expected, the AIDHS/SDS, being an endogamous and relatively homogenous population, was enriched with rare and less common variants. Enrichment of functional variants in cases with HTG along with our focus on individuals with extreme trait (TG) values, increased our power to discover pathogenic variants and aided the discovery of multiple novel/ rare and common/ known variants in splice regions, 5’UTRs, 3’UTRs, intronic, and missense (loss-of-function) variants. Moreover, this ethnic subgroup of Sikhs from North India were enrolled from one single geographic location with shared environmental and cultural traits, which further has reduced the environmental and cultural heterogeneity.

Gene-centric analysis of the identified variants revealed a significant burden of variants for increased HTG risk in *GCKR* (p=2.1×10^−5^), *LPL* (p=1.6×10^−3^) and *MLXIPL* (p=1.6×10^−2^). The GCKR is glucokinase regulatory protein that inhibits glucokinase (GCK) by forming a complex with the enzyme in the liver, which plays a role in glucose homeostasis^44^. Fructose 6-phosphate (F6P) enhances while fructose 1-phosphate (F1P) reduces the GCKR-mediated inhibition of GCK^45^. Lipoprotein lipase (LPL) has long been recognized as an enzyme that hydrolyzes triglyceride-rich lipoproteins to release free fatty acids for energy metabolism^46^. The *MLX1PL* encodes a basic helix-loop-helix leucine zipper transcription factor of the Myc/Max/Mad superfamily. This protein forms a heterodimeric complex and binds and activates carbohydrate response element (ChoRE) motifs in the promoters of triglyceride synthesis genes^47^. Common variants within and around these genes are associated with increased levels of TG and CAD in multiethnic GWAS and metanalysis studies including Sikhs^12, 48^. To test their phenotypic effects and to evaluate metabolic consequences *in vivo,* we focused on three putative variants identified in the human *GCKR* gene by building four transgenic humanized ZF models. These variants were located near the fructose binding site or GCK binding sites at the sugar isomerase domains of the human *GCKR* gene. Evidently, human *GCKR* is about 3 times larger than ZF *GCKR* and it only shows 41% similarity with humans **(Supplementary Figure 2SB)**. Due to the absence of 386 amino acids in the ZF *GCKR* gene, our three functional variants fall outside the ZF GCKR protein.

Despite this dissimilarity, ZF are a well-suited model for studies involving human energy metabolism because the pathways of lipid storage and transport are conserved across species^49^. Further, dietary studies performed in the ZF model for developing atherosclerosis and hepatic steatosis in response to a high-cholesterol diet, revealed the potential strength of this model for analyzing diet-induced phenotypes^50^. In this study, the ZF larvae exposed to a HFD and normal diet, revealed a 2 to 3 fold increase of fat accumulation in hepatocytes in response to the HFD in both TAB-5 and control transgenic fish (normal human *GCKR)* with no apoptosis. However, the observed 4-fold increase in liver fat accumulation with at least one apoptotic cell every hundred hepatic cells, even in the absence of HFD in transgenic *GCKR*^mut^, suggests the impaired function of GCKR due to mutations, which may prevent GCKR from acting promptly in response to the increased concentration of fructose 6-phosphate. This consequently would lead to the uninterrupted release of GCK in the liver to result in the increased uptake of glucose and eventually leading to *de novo* lipogenesis^51^. Alternatively, studies suggest that GCKR stabilizes and protects GCK from degradation. Thus, the increased expression with impaired function of GCKR may result in reduced GCK activity or function, which would give rise to impaired glucose tolerance and hepatic fat accumulation^52^. Evidently, from these studies it appears that the *GCKR* could be a thrifty gene and the functionally disrupted variants in the *GCKR* in Punjabi Sikhs may enhance ectopic fat storage defects even in the absence of HFD, as revealed in the transgenic ZF. Not only did most hepatic cells contain vacuoles of fat, but the structure of hepatocytes was disorganized due to fatty metamorphosis and severe disorganization of the hepatic structure and hepatocyte nuclei with possible steatosis in the absence of HFD in *GCKR*^mut^ compared to *GCKR*^wt^ or WT (TAB-5) **Figure 5C**.

A previously known functional variant (Proline to Leucine) at position 446 of the *GCKR* (rs1260326) was identified as a novel locus for TG metabolism in Caucasian GWAS, and has since been robustly replicated in multiple genome-wide studies of plasma TG^48^. The same variant has also been shown to influence fatty liver disease in children and adults^53^. Of note, the minor risk allele frequency differed significantly between Sikhs (0.27) and other South Asians [e.g. Gujarati Indians (GIH 0.19) and South Asians from Pakistan (ExAC) 0.20], also European Caucasians (0.36). Although our study confirmed the association of this SNP rs1260326 with TG in Sikhs (β 0.09, ± 0.02, p=3.42×10^−5^), the genetic variance explained by this variant was <2% in Sikhs. Whereas, the rs774930016 (representing codon 105) explained 38% of HTG. Both rs760427565 for codon 297 and rs755537970 for codon 553 explained ~25% of HTG genetic variance among carriers. The association of codon 105 with TG remained statistically significant even after controlling for BMI, age, gender, relatedness and T2D **(Table 3)**, suggesting its functional role in increasing HTG risk independent of T2D. Notably, these variants were only restricted to South Asian populations; indeed the rs755537970 (of codon 553) appears to be confined to Punjabi Sikhs **(Table 2).**

We and others have shown that Asian Indian populations may possess a different physiology of obesity ^19, 54–56^. South Asians generally have a non-obese BMI with lower muscle mass and increased visceral fat, which is also associated with their high rates of T2D in the absence of obesity^55, 57–62^. Even results of computed tomography (CT) scans show that Asian Indians have 30% more body fat than age- and BMI-matched African American men, and 21% more body fat than Swedish men^63–65^. Thus, Asian Indians are metabolically obese despite a non-obese BMI. The uneven distribution of fat in insulin sensitive organs like the liver or pancreas increases the risk for development of insulin resistance, T2D, and non-alcoholic fatty liver disease (NAFLD)^**66**^, which are common in Indians. Based on the results of this study, carriers of these evolutionarily conserved variants (specifically S105N and R553W), will have a high risk of ectopic fat deposition and an increased risk for NAFLD in the absence of overt obesity.

Overall, successful humanized transgenic *GCKR* ^*mut*^-expressing *D. rerio* has provided a platform for our ongoing studies to define the precise mechanisms of metabolic derangement perhaps by modulating the GCKR-GCK complex leading to HTG and T2D in humans. Limitations of our study include the lack of data on GCK mRNA, GCKR/GCK protein quantification and GCK activity, which may provide more insight on the putative effects of functional mutations on regulation of GCKR, and clarify the effects of exposure of HFD on the humanized ZF. Our results agree with the earlier reports of targeted improvement of GCK activity by liver–specific GCKR inhibition, which may lower the risk of HTG^45^.

In summary, our study for the first time reports causal role of rare disruptive variants in GCKR for increasing serum TG levels independent of T2D in Punjabi Sikhs. These results may also partly support the “non-obese-metabolic obese phenotype” of Asian Indians linked to increased risk for developing cardiovascular diseases.

## Acknowledgements

Authors thank all the participants of AIDHS/SDS who made this study possible. We thank the Stephenson Cancer Center at the University of Oklahoma, Oklahoma City for the use of Histology and Immunohistochemistry Core, which provided Processing, Embedding and Tissue staining service, and Nikon Microscopic imaging service. Authors thank Dr. Amnon and Dr. Schlegel, University of Utah, for providing us the *D. rerio* liver fatty acid binding protein (L-FABP) promoter. Technical help provided by Drs. Bishwa Sapkota, Jayaraman Muralidharan, Anil Singh, Louisa Williams, and Sheeja Aravindan is duly acknowledged.

## Author Contributions

Conceived and designed the experiments: DKS; Data analysis, SNP genotyping for replication analysis, bioinformatics, plasmid cloning using Tol2 technology RH, CT, CB, HM, PW; Provided Fish breeding and care MWM; advice and help in Tol2 technology JKF; Immunohistochemistry and slide reading KAF and CVR and SL; DKS wrote and JKF helped in manuscript editing; All authors read and approved the final manuscript.

## Supplementary Information (Legends)

**Supplementary Figure 1S.** Flow Chart summarizes research design, targeted sequencing, replication and functional studies workflow.

**Supplementary Figure 2S A-B.** GCKR protein alignment of human vs. zebrafish using BLAST (https://blast.ncbi.nlm.nih.gov/Blast.cgi)

**(Figure 2S-A).** Upper gray horizontal bar is the protein sequence of human GCKR (NP_001477.2) with 625 amino acids, and the lower red horizontal block is the zebrafish GCKR (XP_002665191.3) protein sequence.

**(Figure 2S-B).** Protein sequence alignment of human and zebrafish GCKR. Upper rows represent human residues and lower rows represent zebrafish residues starting at codon 387 of human GCKR. The black spaces and + symbols indicate low degree of homolog between human and zebrafish, only 94 out of 233 (41%) residues showed complete alignment.

**Supplementary Figure 3S.** Additional supplemental figures of WT and transgenic Zebrafish livers at 40X magnification.

**Supplementary Figure 4S.** Additional supplemental figures of WT and transgenic Zebrafish livers at 4X magnification.

**Supplementary Table 1S:** Gene-centric association of coding variants using combined multivariate and collapsing (CMC) and SKAT-O (uniform) analyses.

**Supplementary Table 2S:** Summary of classification and distribution of 2361 high quality variants identified in the discovery cohort using targeted sequencing.

**Supplementary Table 3S: (Table 3S-A).** Demographic and clinical traits of carriers of GCKR S105N (rs774930016) variant in subjects from the AIDHS. **_(Table 3S-B)**. Demographic and clinical traits of carriers of GCKR R297Q (rs760427565) variant in subjects from the AIDHS. **(Table 3S-C).** Demographic and clinical traits of carriers of GCKR R553W (rs755537970) variant in AIDHS.

